# Cost-efficient high throughput capture of museum arthropod specimen DNA using PCR-generated baits

**DOI:** 10.1101/333799

**Authors:** Alexander Knyshov, Eric R.L. Gordon, Christiane Weirauch

**Author notes:** corresponding author email and ORCID, orcid.org/0000-0002-2141-9447. Current affiliation: University of Connecticut, Ecology and Evolutionary Biology, Storrs, CT, USA.

## Abstract

1. Gathering genetic data for rare species is one of the biggest remaining obstacles in modern phylogenetics, particularly for megadiverse groups such as arthropods. Next generation sequencing techniques allow for sequencing of short DNA fragments contained in preserved specimens >20 years old, but approaches such as whole genome sequencing are often too expensive for projects including many taxa. Several methods of reduced representation sequencing have been proposed that lower the cost of sequencing per specimen, but many remain costly because they involve synthesizing nucleotide probes and target hundreds of loci. These datasets are also frequently unique for each project and thus generally incompatible with other similar datasets.
2. Here, we explore utilization of in-house generated DNA baits to capture commonly utilized mitochondrial and ribosomal DNA loci from insect museum specimens of various age and preservation types without the a priori need to know the sequence of the target loci. Both within species and cross-species capture are explored, on preserved specimens ranging in age from one to 54 years old.
3. We found most samples produced sufficient amounts of data to assemble the nuclear ribosomal rRNA genes and near complete mitochondrial genomes and produce well-resolved phylogenies in line with expected results. The dataset obtained can be straightforwardly combined with the large cache of existing Sanger-sequencing-generated data built up over the past 30 years and targeted loci can be easily modified to those commonly used in different taxa. Furthermore, the protocol we describe allows for inexpensive data generation (as low as ∼$35/sample), of at least 20 kilobases per specimen, for specimens at least as old as ∼1965, and can be easily conducted in most laboratories.
4. If widely applied, this technique will accelerate the accurate resolution of the Tree of Life especially on non-model organisms with limited existing genomic resources.

## Introduction

Natural history museums host troves of biological material and sometimes the only known representatives of extinct or rare species (Coddington, Agnarsson, Miller, Kuntner, & Hormiga, 2009; Lim, Balke, & Meier, 2011). In these cases, museum specimens represent the only accessible sources of genetic data for a given species and gathering data from such specimens in a cost-effective way is one of the primary obstacles yet to be overcome in modern phylogenetics. Specimens in museums may also allow for the inclusion of a temporal variable into analyses by comparing DNA sequence of individuals across different sampling dates and can even be used for the analysis of short-term evolutionary trends (Hartley et al., 2006; DiEuliis, Johnson, Morse, & Schindel, 2016).

Preservation conditions of museum material can dramatically impact the viability of obtaining DNA sequence data. Traditional approaches used amplification of target regions of DNA followed by Sanger sequencing. This method is highly dependent on residual DNA fragment size and the proportion of endogenous DNA in the extract. While targeting shorter gene regions can mitigate the issue of DNA fragmentation, a low endogenous content is harder to overcome (Burrell, Disotell, & Bergey, 2015), and even innovative new PCR techniques are only capable of somewhat reliably amplifying fragments of less than 600 bp (Mitchell, 2015). The development of next generation sequencing (NGS) has expanded the array of methods for DNA sequencing from museum specimens. For whole genome sequencing, an NGS library is prepared from the original DNA extract and this library is then combined with other samples for multiplex sequencing and allocated a certain proportion of reads on a sequencer, depending on desired sequencing depth and the total budget (Cridland, Ramirez, Dean, Sciligo, & Tsutsui, 2018; Kanda, Pflug, Sproul, Dasenko, & Maddison, 2015; Maddison & Cooper, 2014). However, even low-coverage whole genome sequencing is currently still prohibitively expensive for all but very well-funded projects or studies focusing on relatively few samples.

As a way to decrease the cost per sample while still generating sufficient amounts of data for accurate phylogenetic placement, several methods of reduced representation sequencing have been proposed. Typically, these methods include selective hybrid capture of target loci, where the type and number of loci being captured depends on the scope and context of the study. During the past few years, utilization of commercially synthesized probes or microarray kits for the capture of conserved DNA regions has become popular for phylogenetic studies and can be applied to historical museum specimens (Bi et al., 2013; Blaimer, Lloyd, Guillory, & Brady, 2016; McCormack, Tsai, & Faircloth, 2015). These kits are designed based on existing reference genomes or transcriptomes and typically enrich many loci (∼500-5000), thus a large amount of data is generated for each sample, but the cost per sample is relatively high. Other kits are designed to enrich mitochondrial genomes, including a kit specifically designed for mitochondrial DNA across insects (Liu et al., 2016). However, all methods relying on commercially synthesized kits are relatively expensive and might not be feasible for low-budget projects. These kits are also limited by the original design and probe composition cannot be adjusted after synthesis.

These limitations led us to explore an approach that uses in-house generated DNA baits for hybrid enrichment (Maricic, Whitten, & Pääbo, 2010). These baits can be produced from amplicons generated by PCR of short gene regions (Peñalba et al., 2014), or by long-range PCR of complete mitochondrial (Li et al., 2015; Maricic et al., 2010) or chloroplast genomes (Mariac et al., 2014), or even from ddRAD library fragments (Suchan et al., 2016). PCR-generated baits have so far only been applied to vertebrates and plants, and only in a few cases tested on archival specimens (Li et al., 2015). Appealing features of this approach include affordable synthesis of baits, independence from the need of a good quality reference, and flexibility of the synthesis workflow for low-cost modifications of the bait set (e.g., pooling different combinations of bait amplicons, using same primers to obtain bait amplicons from different taxa, or generating additional baits with new sets of primers).

The diversity of arthropods is staggering, with estimates of about 80% of species still undescribed (Stork, 2018) and an increasing number of species going extinct every day (Hallmann et al., 2017). While modern phylogenetic studies of vertebrates sometimes approach complete sampling of extant diversity, complete extant sampling of any large clade of arthropods is almost impossible due to the abundance of rare species, limited material, and the huge diversity of arthropods (Coddington et al., 2009; Lim et al., 2011). However, near-complete sampling is useful for many downstream analyses, including unbiased estimation of lineage diversification rates (Cusimano, & Renner, 2010; Cusimano, Stadler, & Renner, 2012; Höhna, Stadler, Ronquist, & Britton, 2011). Scientists have just started to utilize the enormous resources of arthropod specimens deposited in natural history collections for gathering large DNA datasets (Stork, McBroom, Gely, & Hamilton, 2015; Stork, 2018). We argue that insects in particular are an apt test case for the application of new NGS approaches to illuminating the dark areas in the Tree of Life, because most material in entomological collections is stored as dried and pinned or point-mounted specimens, which are often suitable for the retrieval of fragmented DNA. Previous applications of this approach on vertebrate and plant samples employed destructive extraction protocols to generate adequate amounts of DNA for capture. But DNA extraction can be performed without destroying external or genitalic morphological features and from individual and small specimens as in many insects. For a complete taxonomic sampling of large clades, already existing data should be compatible with character-rich new datasets generated at low costs.

Here, we test the efficiency of PCR-generated DNA baits (targeting the mitochondrial genome, nuclear ribosomal operon, and one nuclear protein-coding gene) to capture DNA sequences from museum-deposited insect specimens with different collection dates, preservation methods, and evolutionary relatedness, using phyline plant bugs (Insecta: Hemiptera: Miridae: Phylinae) as our test case. These loci were selected for optimal integration with existing, Sanger-based sequence data and to allow adequate coverage when multiplexing hundreds of libraries. Plant bugs are a group of > 11,000 described species that include serious plant pests and beneficial insects (Cassis & Schuh, 2012). Phylogenetic hypotheses for the entire group are in their infancy (Jung & Lee, 2011), but studies targeting selected subfamilies including the Phylinae now provide testable hypotheses (Konstantinov & Knyshov, 2015; Menard, Schuh, & Woolley, 2014; Namyatova, Konstantinov, & Cassis, 2015; Tatarnic & Cassis, 2012). The taxonomic diversity of plant bugs in the Western U.S. is fairly well understood (Cassis & Schuh, 2012; Weirauch et al., 2016), but few species have been incorporated into phylogenetic analyses, and some are only known from the type specimen(s). As the first test case, we selected a putatively monophyletic group of native oak-associated plant bugs, the so called “Orange Oak Bugs” (OOB) (Weirauch, 2006a, 2006b), where some species may be monophagous on specific species of oaks, while at least two widespread and polymorphic species (*Phallospinophylus setosus* Weirauch and *Pygovepres vaccinicola* (Knight)) feed on a variety of host plants (including Fagaceae, Rhamnaceae, and Rosaceae). We sampled specimens of these two species from a range of localities and host plants, together with several additional species of OOB that had not yet been included in phylogenetic analyses (Menard et al., 2014) to test efficacy of capture across closely related samples and to investigate potential cryptic host plant races. As second test case, we selected the genus *Tuxedo* Schuh with seven described species associated with host plants in several families (Schuh, 2004); phylogenetic relationships within this genus are unknown. We aimed to sample several individuals from each of the seven species, including paratype specimens, to investigate capture efficiency at deeper phylogenetic levels and to explore host plant shifts within the genus. Both datasets were analyzed together with a Sanger-derived phylogenetic dataset of Phylinae (Menard et al., 2014), demonstrating the feasibility of combining existing and newly generated NGS data.

## Material and methods

### Taxon Sampling and Vouchering

Specimens for this study were loaned from the American Museum of Natural History (AMNH), the Entomology Research Museum (UCRC), and the Zoological Institute, Russian Academy of Sciences (ZISP). Tentative voucher identification was done based on habitus and host association data using Weirauch (2006a, 2006b) and Schuh (2004). Age of specimens at the moment of DNA extraction varied from one to 54 years. Specimens of *Tuxedo, Leucophoroptera* Poppius, *Ausejanus* Menard and Schuh, and *Pseudophylus* Yasunaga were imaged using a Leica DFC 450 C imaging system. Image vouchers and specimen information are available through the Heteroptera Species Pages (http://research.amnh.org/pbi/heteropteraspeciespage/). After clearing soft abdominal tissues during the DNA extraction process, we examined male genitalic characters to confirm our tentative identifications. In cases where different diagnostic characters were in conflict (e.g., in some *Tuxedo* spp., see results and discussion), we based our identification on genitalic characters.

### DNA Extraction

In most cases, only the abdomen (1-1.5 mm in length) was used for non-destructive DNA extraction, which was performed using a Qiagen DNeasy® Blood & Tissue kit (for relatively fresh ethanol specimens) or a combination of the previous kit with a Qiagen QIAquick® PCR purification kit (for dry specimens, see supplemental text S1), since the latter is commonly used for DNA extraction from degraded samples (Lee et al., 2010; Yang, Eng, Waye, Dudar, & Saunders, 1998). Abdomens were soaked in the extraction buffer, such that cuticular structures remain undamaged, and mirrored standard dissection procedures for plant bug specimens. This approach allows for subsequent remounting of the abdominal cuticle and genitalia with the rest of the specimen, or in a genitalic vial.

### Bait Synthesis

Freshly collected specimens of *Phallospinophylus setosus* and *Tuxedo drakei* Schuh were selected as bait donors for the OOB and *Tuxedo* subprojects, respectively. Primers for obtaining long range PCR products are listed in Table S1. Details on the primer design are available in supplemental text S1. Target regions included mitochondrion, nuclear ribosomal operon, and a fragment of the cytoplasmic dynein heavy chain gene.

To prepare baits, six long-range (LR) PCRs per specimen were performed. For this and all subsequent PCR described in this paper, we used Takara PrimeSTAR® GXL polymerase, a hot-start high-fidelity enzyme that is able to amplify long products. The PCR mix contained 10 µl PrimeSTAR® GXL buffer, 4 µl 2.5M dNTPs, 1 µl PrimeSTAR® GXL polymerase, 32 µl water, 1.5 µl of each primer (10 µM), and 1 µl of DNA template. The thermocycler program included initial denaturation at 98° for 3 min, 35 cycles of denaturation for 10 sec at 98°, followed by annealing at variable temperatures for 15 sec, followed by elongation at 68° for a variable amount time, and with the final incubation at 68° for 15 min. Additional details on long-range PCR conditions are available in Table S1.

After clean up with custom Solid Phase Reversible Immobilization (SPRI) beads (Glenn et al., 2016; Rohland, & Reich, 2012), mitochondrial, nuclear ribosomal, and nuclear protein-coding products were mixed in molar ratios of 1:1:5, following recommendations of Peñalba et al. (2014) regarding capture of low copy nuclear genes. Mixtures were diluted to the volume of 100 µl and sonicated on a Diagenode Bioruptor® UCD-200 with 30/30 cycles for 6 runs of 5 minutes. Sheared PCR products were subjected to a bait library preparation generally following the protocol of Li et al. (2015) with the exception that regular dNTPs instead of a dUTP-containing mixture were used, since NaOH melting was used to subsequently elute captured libraries instead of off-bead amplification. Three pools of ready-to-use bait were produced by amplifying M13-adaptor-ligated bait libraries with 5’ biotinylated primers using PCR conditions outlined above with the following modifications: 6 µl of template was used, and annealing temperature set to 55°.

### Preparation of Illumina-compatible Libraries

Since DNA sequence of bait donors was also of interest in this project, we also sequenced amplicons used for bait production. These LR PCR products were mixed in equimolar ratios and sonicated as described above. Following sonication, Illumina®-compatible libraries were prepared using the protocol from Li, Hofreiter, Straube, Corrigan, and Naylor (2013), with the following modifications: end prep mix contained 50% 2X Takara EmeraldAmp® GT PCR mix and after incubation at 25° for 15 min and 12° for 5 min was incubated at 72° for 20 min in order to obtain a-tailed fragments. We utilized with-bead SPRI method as originally described in Fisher et al. (2011), carrying same SPRI beads through the library preparation steps. T-tailed loop adaptors from NEBNext® Multiplex Oligos for Illumina® kit E7600s were ligated to the DNA and a PCR with indexing primers from the same kit was conducted using PCR conditions outlined above with the following modifications: 6 µl of template was used, annealing temperature set to 60°, number of cycles set to 16.

To prepare target libraries, DNA extracts were run on a gel with Biotium GelRed® premixed loading buffer in ratios 1:2 to check average fragment size and determine if sonication was needed (i.e., for younger samples). These DNA extracts were quantified using Qubit™ fluorometer, and for more consistent sonication results approximately 70 ng of DNA (where possible, also see Table 1) were used for sonication. Library preparation followed the protocol outlined above with the exception that after adaptor ligation, libraries were amplified with short IS7/IS8 primers following Li et al. (2013). The same PCR conditions as above were used, however number of cycles were varied from 16 to 21 depending on the amount of starting material.

**Table 1.**
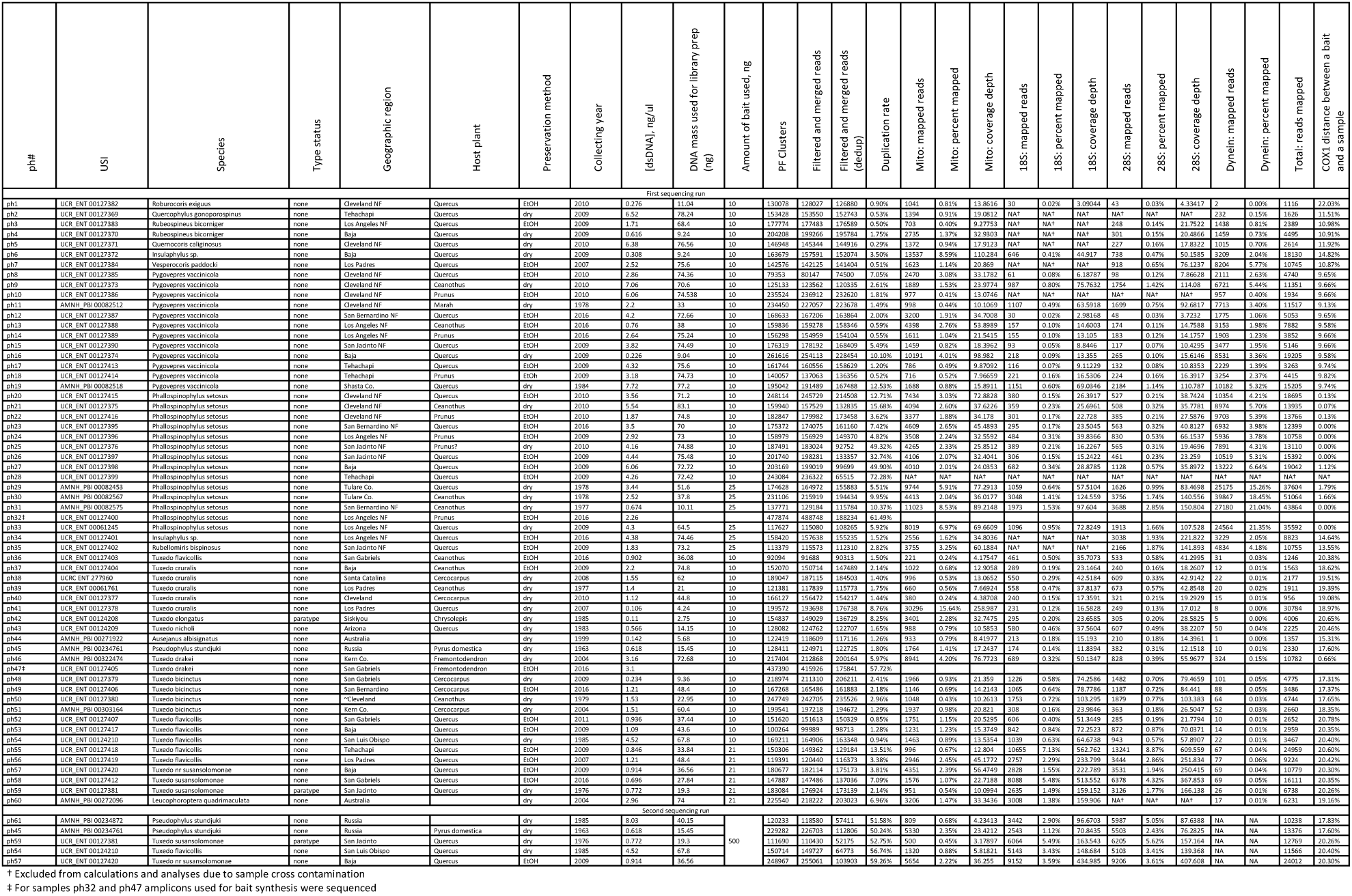
List of samples used in the project, voucher specimen information, and sequencing information.

### First Sequencing Run – Target Capture, Pooling and Sequencing

In our first sequencing run, target captures generally followed the protocol of Li et al. (2015). Every sample was captured individually as in Li et al. (2015), 10 µl of Invitrogen Dynabeads® M-270 and 10 ng of bait library was used for most samples, whereas all remaining bait library was used for the last few captured samples (for details on bait amount used, see Table 1). DNA concentration of input target library was not quantified, and we used 6 µl of target library in each capture reaction. Elution was conducted with NaOH melting as in Maricic et al. (2010), and double capture was performed following suggestions of Peñalba et al. (2014). After the second round of capture, the supernatant was cleaned, and eluted in 50 µl of 10 mM Tris-HCl. Post-capture PCR followed the same PCR procedure as outlined above, however indexing primers and 20 µl of template were used, and variable number of cycles was performed (16-24).

After indexing PCR, products were cleaned and normalized with Just-a-Plate™ 96 PCR Purification and Normalization Kit. Since using Bioanalyzer on all 60 samples was prohibitively expensive, libraries were first run on a gel with GelRed® to check average fragment size, pooled together into nine groups according to their size, which were then analyzed on a single Bioanalyzer chip to obtain more accurate fragment size distribution. Then libraries were pooled equimolarly with the exception of sheared amplicon libraries (samples ph32 and ph47), which were pooled at twice higher concentrations. The pool of 60 indexed libraries then was mixed in molar proportion of 50:50 with unrelated samples from other projects and sequenced on a single run of Illumina® MiSeq® V3 2×300bp at the UCR IIGB Core Facility.

### Second Sequencing Run – Library Preparation, Target Capture and Sequencing

In the second sequencing run, we followed the protocol of Maricic et al. (2010) with modifications. DNA extracts from the same specimen of *Tuxedo drakei* as above was used as a source for bait preparation. The procedure differed from described above in that only nuclear rRNA operon and mitochondrial PCR products were used. We extracted one more specimen of *Pseudophylus* and prepared a library as outlined above. Five libraries (samples ph45, ph54, ph57, ph59, and a new *Pseudophylus* library) were carried through indexing PCR, quantified using Qubit™, checked on an agarose gel, and pooled equimolarly to obtain about 450 ng of DNA. Because indexed libraries were used, we added additional blocking oligos as in Maricic et al. (2010) to block longer adaptor fragments. Approximately 500 ng of bait and 5 µl of Dynabeads® as in Maricic et al. (2010) were used for each round of capture (two rounds total as in the first sequencing run). Post-capture amplification was done using IS5 and IS6 primers and was carried over in two aliquots. After PCR, the products were combined and purified, they were then sequenced on 5% of another Illumina® MiSeq® V3 2×300bp run at the UCR IIGB Core Facility.

### Post-Sequencing Data Processing

Raw sequences were demultiplexed and adaptors were removed using bcl2fastq software (Illumina®) at the UCR IIGB Core Facility. Trimmomatic v0.36 (Bolger, Lohse, & Usadel, 2014) was used to trim off low quality ends of the sequences as well as perform more thorough adaptor trimming. Reads were assembled into contigs with SPAdes (Bankevich et al., 2012). In cases where assembly did not yield complete target regions, we obtained them by mapping shorter contigs onto full length assemblies of other related samples. Assembled contigs were checked for misassembled regions and manually curated in Geneious v.10 (https://www.geneious.com, Kearse et al., 2012). We mapped reads on these contigs using BWA (Li & Durbin, 2009) to assess the coverage depth (see Table 1), prior to average coverage calculations, reads were deduplicated using PRINSEQ (Schmieder & Edwards, 2011).

We aligned all resulting 18S, 28S, and mitochondrial contigs using MAFFT v.7 (Katoh & Standley, 2013). Manual inspection of alignments and trimming was performed. Since accurate assembly of the mitochondrial control region with short reads without a close reference was problematic due to presence of repeats, we excluded it from the analysis. The remainder of the mitochondrion was annotated by aligning it with mitochondrial genome of another plant bug available on GenBank (NC_024641.1).

### Phylogenetic Analysis

For phylogenetic analysis, the dataset was concatenated and divided into 18 partitions with protein coding genes split further into codon positions. Substitution models and partitioning scheme were optimized using PartitionFinder 2.1.1 (Lanfear, Frandsen, Wright, Senfeld, & Calcott, 2016) or ModelFinder (Kalyaanamoorthy, Minh, Wong, von Haeseler, & Jermiin, 2017), which is an IQ-TREE built-in model and partition test. Phylogeny estimation was performed in RAxML v8.2.11 (Stamatakis, 2014) and IQ-TREE v1.5.4 (Nguyen, Schmidt, von Haeseler, & Minh, 2014). Branch support was calculated using Rapid Bootstrap (Stamatakis, Hoover, & Rougemont, 2008) which is shown on Fig. 2, Ultrafast Bootstrap (Minh, Nguyen, & von Haeseler, 2013), and SH-aLRT (Anisimova & Gascuel, 2006), which are shown on Figs S2 and S3.

**Figure 1.**
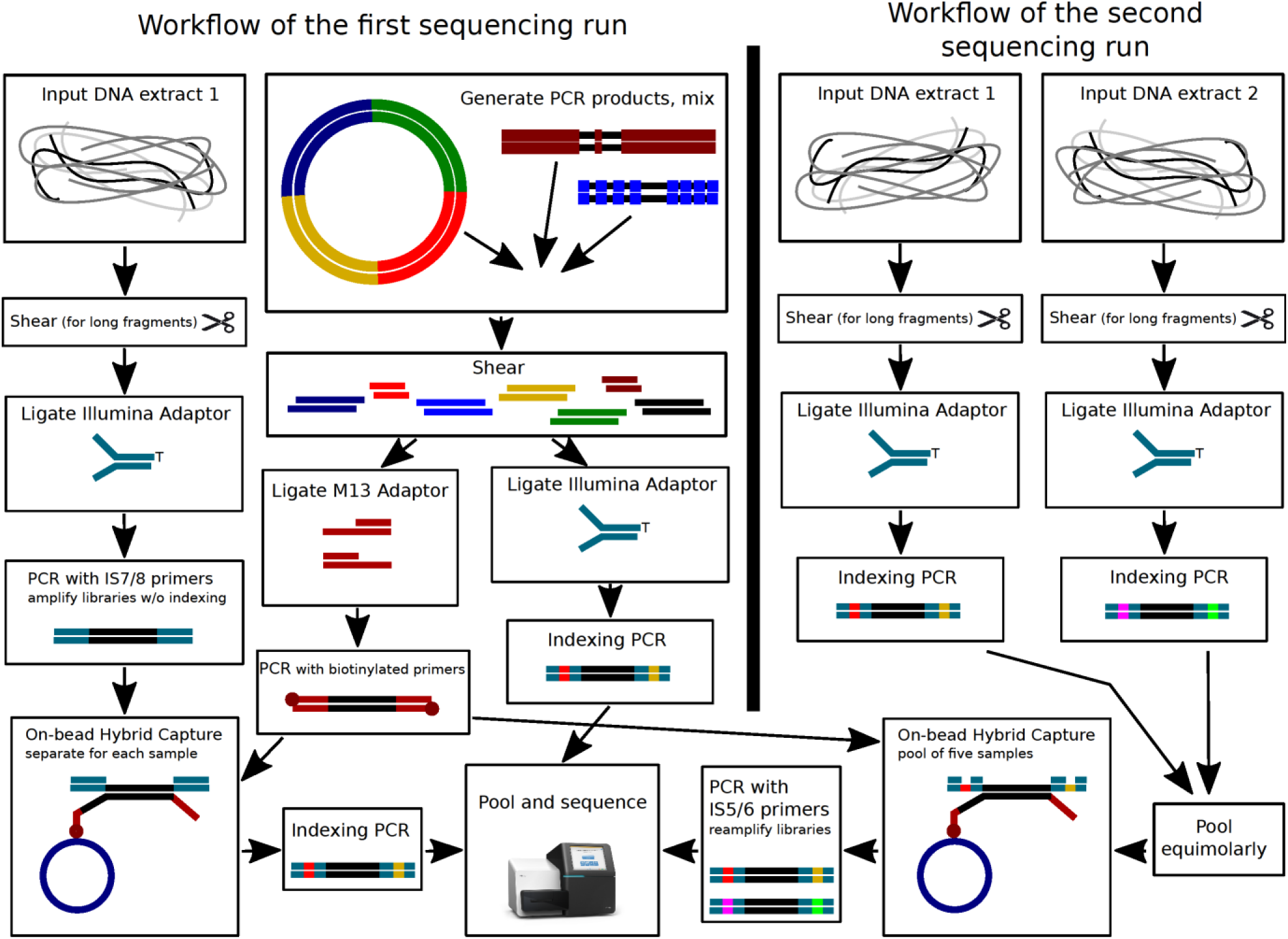
Procedure flowchart.

**Figure 2.**
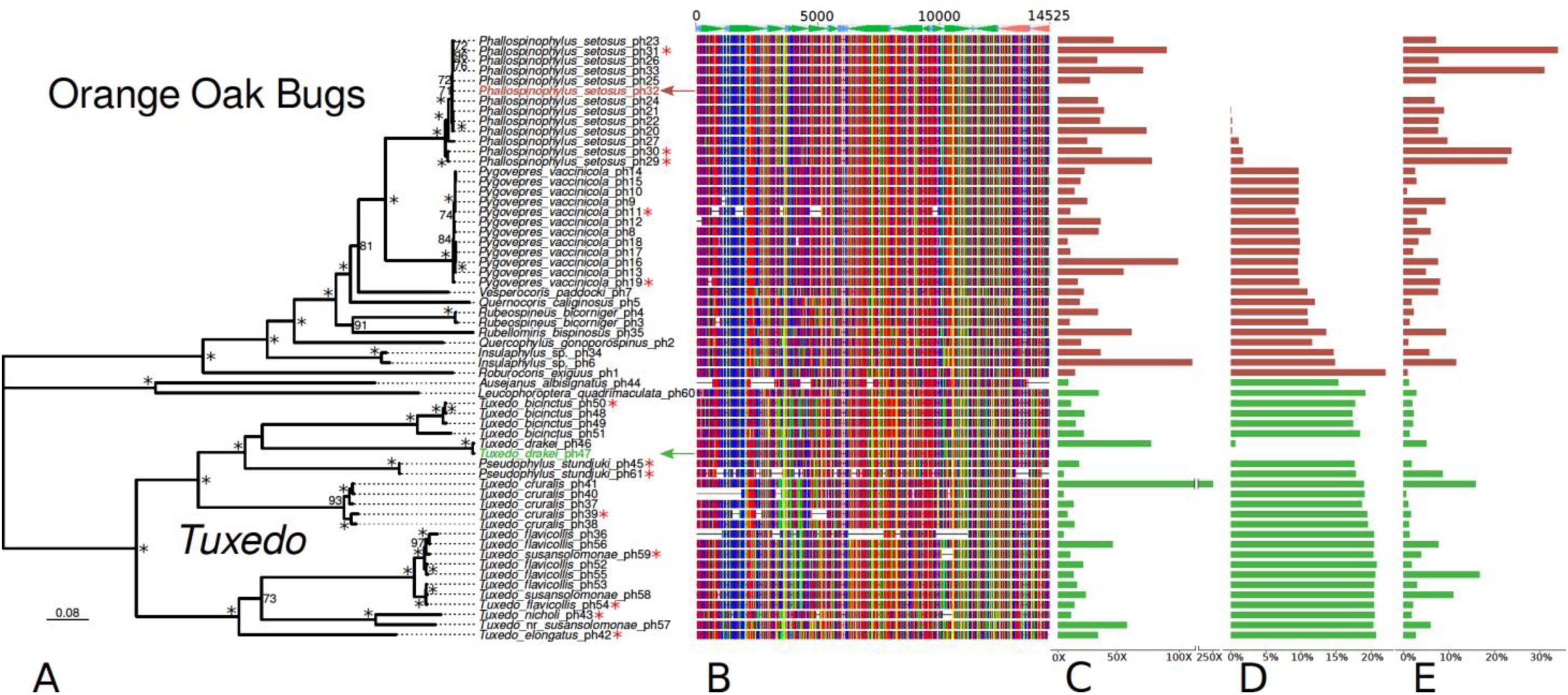
A. Combined phylogeny of the OOB and *Tuxedo* subprojects, generated in RAxML, values at nodes represent Rapid Bootstrap Support, values below 70 are not shown, asterisks indicate full support, arrows denote bait samples for the OOB (red) and *Tuxedo* (green) subprojects, red asterisks denote samples older than 20 years. B. Mitochondrial alignment completeness, control region excluded. C. Average coverage of mitochondrial contig(s), control region excluded. D. Pairwise COX1 distances between a bait and a captured sample. E. Total percent of reads mapping to target including mitochondrial genome, 18S, 28S, and dynein.

To test how well our data can be combined with previously generated data, we combined our data with the dataset of Menard et al. (2014) which is the most comprehensive set of genetic data for related species. After downloading the sequences from GenBank, we extracted only 18S, 28S, 16S and COX1 sequences from our data, performed alignment and manual trimming. Alignments were then concatenated, optimized for model and partitioning scheme, and phylogenetically analyzed as above.

Illustrations for Figs 2, 3, and S1 were drafted using R v3.4.3 and packages APE (Paradis, Claude, & Strimmer, 2004), phytools (Revell, 2012), ggtree (Yu, Smith, Zhu, Guan, & Lam, 2016) and ggbio (Yin, Cook, & Lawrence, 2012). Relief image for Fig. 3 was taken from the SimpleMappr website (http://www.simplemappr.net/).

**Figure 3.**
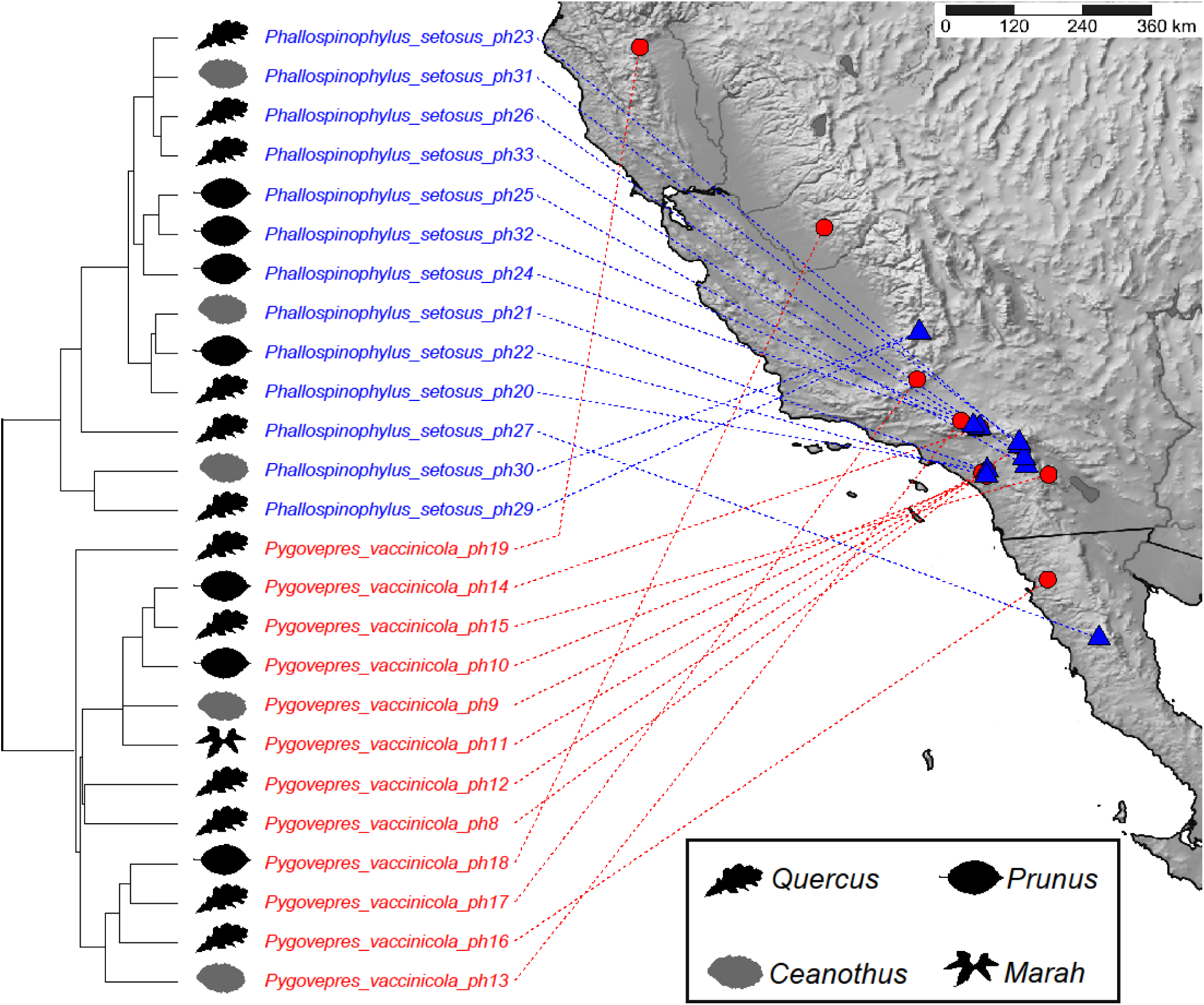
Host and distribution data for the Orange Oak Bug subproject, aligned with phylogeny (branches not to scale) and with host plant of specimens mapped using representative leaf shapes of plant genus.

## Results and Discussion

### Expenses

Total expenses after bait and target library preparations, target capture, and sequencing and including all reagents and supplies came to about $54 per specimen or about $2.8 per 1 Kb of data in the first sequencing run, and about $39 per specimen or about $2.1 per 1 Kb of data in the second sequencing run (Table S2). Our estimates suggest that pooled capture together with using a higher throughput sequencer (e.g., a HiSeq® lane or a NextSeq® run) can generate the same amount of data for about half the price (up to $25 per specimen), however a greater number of samples (at least 360) need to be pooled together to efficiently utilize the sequencer.

### DNA extraction

The amount of DNA extracted greatly varied across samples (see Table 1). The minimum amount of DNA that was used for library preparation was 2.75 ng (sample 42). The average fragment size for ethanol preserved material was large: we always detected a bright band larger than 10 Kb in size, with many extracts also with a smear of fragments spanning down to 300 bp. For dry point-mounted material we observed two types of fragmentation: extracts that had fragments of 500-700 bp on average in addition to long (∼8Kb) fragments (dry specimens collected within past ten years), and extracts with only fragments shorter than 1000 bp (dry specimens collected more than ten years ago).

### Sequencing and assembly

A total of 45% of an Illumina® MiSeq® V3 lane was used for the samples in the first sequencing run. The amount of reads obtained per sample is listed in Table 1 (average of 152670, σ = 39070). For bait samples, we obtained full bait contigs for the nuclear ribosomal operon, the dynein fragment, and entire mitochondrial genome, although unambiguous assembly of the control region was problematic due to lack of a close reference and long read data. For other samples, we obtained full or partial mitochondrial contigs and nuclear ribosomal gene contigs for the majority of samples (see Table 1). Mitochondrial completeness is indicated on Fig. 2B and excludes the control region, and mitochondrial average coverage depth is indicated in Fig. 2C. We were able to obtain reliable ribosomal data for 48 taxa, but some sequences exhibited cross contamination of about 1% of reads by the bait taxon as detailed in supplemental text S1. In the second sequencing run, we observed a higher percent of ribosomal operon reads on target (on average 7.33% in the second run compared to 2.1% in the first run for the same samples) and both for recaptured libraries, as well as for the library prepared after the first sequencing run was complete (ph61), we have not detected contaminating reads.

### Capture efficiency

Percent of reads on target varied from 0.61% to 33.95% and was on average 8.19% in the OOB subproject and 4.02% in *Tuxedo* subproject. The percent of reads on target was slightly larger for samples that are close to bait specimens (Figs 2A, 2E, Fig. S4). We also observed a significant variation of percentage on-target across samples of close phylogenetic relatedness, which may be attributed to variation in total amount of target DNA submitted to capture reactions (equal volumes of target libraries were used in all reactions). Capture in the *Tuxedo* subproject performed worse, which could be attributed to the higher sequence divergence from the bait (Fig. 2D).

Baits for a nuclear protein-coding gene (dynein) performed unsatisfactorily, even though they were five times more concentrated. Although we do not have a clear explanation as to why this bait performed suboptimally (see supplemental text S1), the large middle intron may have been detrimental for bait efficacy.

On the contrary, mitochondrial baits were only 14.3% of total bait pool, yet were able to considerably enrich for mitochondrial DNA. Typical sequencing of non-enriched DNA libraries from insect museum specimens yields from 0.002% to 0.08% of total reads mapping to the mitochondrial genome (Staats et al., 2013), however, we recovered on average of 2.13% (σ = 2.59%, range 0.24%-15.64%) representing an enrichment of at least 25x on average for our first sequencing run. Given the amount of reads we allocated for our samples, an unenriched library would produce only about 120 mitochondrial reads, where we achieved on average ∼3,500 reads (an enrichment of 29x), sufficient for assembling the whole mitochondrial genome.

Suboptimal capture performance in our first sequencing run could be also attributed to the amount of bait. Overall, we observed an increase in the amount of reads on target in capture reactions where more bait was used (Table 1, samples ph29-ph31, ph33-ph35, and ph55-ph60). Thus, we repeated sequencing of five selected samples captured with a modified protocol (see Materials & Methods) where more bait was used. We also explored a pooled capture approach, which is significantly cheaper than the individual sample captures. In the result of the second sequencing run, we observed on average 8.65% on target reads as opposed to 3.42% for the same samples in the first sequencing run (see Table 1). We also noticed a larger variation of total amount of reads received for a given sample in the pool. This might be due to unequal divergence of samples in the pool with the respect to the bait or difference in library quality due to the age of the specimens. Because of this, we recommend balancing sample pools prior to capture and performing individual captures for sensitive samples.

Our results show no difference in capture efficiency as related to the age of the specimen (Fig. 2, specimens older than 20 years denoted with red asterisks). We thus expect that even older specimens can be used (Blaimer et al., 2016), but for this pilot study the youngest available specimens were chosen. Further adjustments of hybridization temperature and duration may further improve capture success, however need to be modified on an individual basis.

### Phylogenetic analyses

Using the obtained data, we reconstructed a well resolved phylogeny, contributing new insights. Our phylogenetic analysis supports the monophyly of *Tuxedo* + *Pseudophylus*, the OOB clade, *Phallospinophylus setosus* and *Pygovepres vaccinicola* with the highest branch support (Fig. 2A, Fig. S2). As part of the *Tuxedo* subproject, we sampled two specimens of *Pseudophylus stundjuki* (Kulik) since this species from Far East Asia rendered the Western Nearctic *Tuxedo* paraphyletic in a previous analysis (Menard et al., 2014). “*Tuxedo*” is here confirmed to be paraphyletic with respect to *Pseudophylus*, after thorough examination of our sequence data and comparison with data from Menard et al. (2014) and Jung and Lee (2011). All primarily Fagaceae-feeding species of “*Tuxedo*” form a well-supported monophyletic group. Species other than *Tuxedo flavicollis* (Knight) and *Tuxedo susansolomonae* Schuh were recovered as monophyletic and conform with genitalic-based identifications. Phylogenetic analysis recovered two highly supported monophyletic groups within the *T. flavicollis/susansolomonae* species group, however composition of each group is not congruent with either genitalic structure or coloration. One specimen (ph57) initially identified as *T. susansolomonae* is distantly related from other members of *T. flavicollis/susansolomonae* clade and is recovered as sister taxon to *T. nicholi* (Knight), and likely represents an undescribed species. Species within the OOB clade represented by multiple specimens are monophyletic with high support. Our analysis did not find support for our hypothesis on the presence of host plant races within each of the widespread and polyphagous OOB species (Fig. 3). In contrast, the phylogenetic structure in *Phallospinophylus setosus* is more likely explained by geographic proximity between sampled localities.

Combined with existing data of Menard et al. (2014), phylogenetic hypotheses inferred from our dataset are congruent with those presented in prior studies (Fig. 4, Fig. S3). Deep level relationships within Oncotylina as well as the monophyly of the subtribe itself remain poorly supported based on this data set. As in Menard et al. (2014), “*Tuxedo”* + *Pseudophylus* are recovered as the sister group to Leucophoropterini, although with low support. Sampled species of Leucophoropterini were recovered in expected phylogenetic positions.

**Figure 4.**
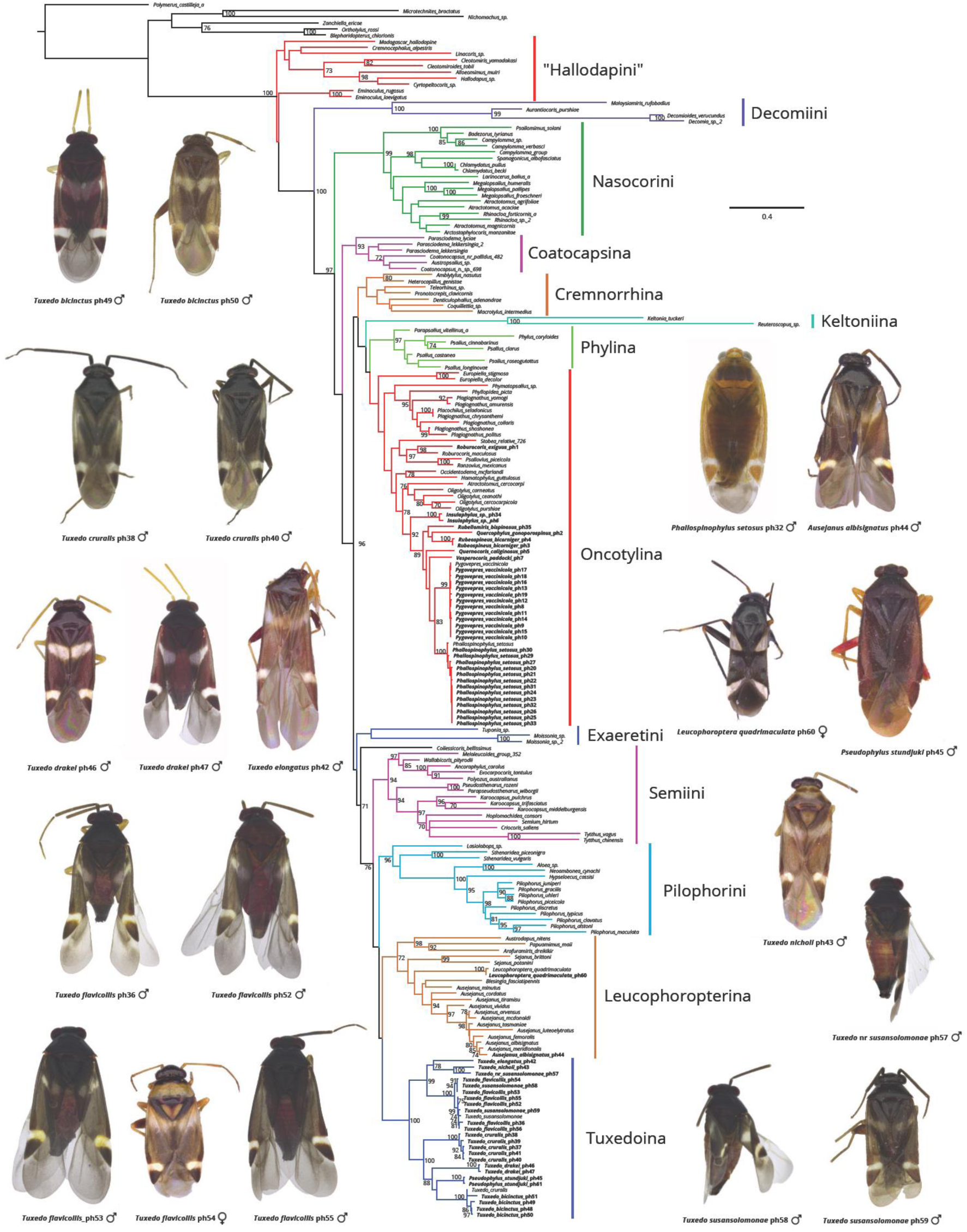
Phylogeny of Phylinae, generated in RAxML, with specimens for which new data was gathered in bold font, values at nodes represent Rapid Bootstrap Support, values below 70 not shown.

## Conclusions

In conclusion, we were able to cost-efficiently ($2.8/sample/Kb) sequence long-range PCR products as well as perform hybrid enrichment using in-house generated baits and obtain DNA sequences (∼20 Kb) from archival specimens (up to 54 years old) using a minimal amount of DNA. This approach offers a much lower cost of bait production than other approaches, however, especially if LR PCR is chosen for amplicon generation, a high-quality sample of a related species is needed. While it is hard to scale up this method to produce baits for 500 targets, it is well suited to generate commonly used high-copy gene sequences for both archival and recently collected samples. It fits within a narrow ‘Goldilocks’ zone in terms of adequate data for accurately reconstructing phylogenies and relative cost effectiveness with the ability to multiplex at least ∼120 individuals per MiSeq® run given the number of loci captured. While the amount of reads on target in our project was not high, we were able to assemble genes of interest for most captured samples.

Data obtained showed no evidence for host plant races in OOB. For *Pygovepres*, we could not detect any phylogenetic structure within the species, whereas the structure within *Phallospinophylus* could be explained by distribution. We also reconstructed the phylogeny of the genus *Tuxedo* and sampled all described species, some of which were rarely collected species that are based on specimens from type series.

Finally, it is straightforward to combine such data with previously generated data using conventional Sanger sequencing. Commonly used primers for different genes for use in phylogenetic analysis of other groups are easy to add to our protocol. When applied to museum specimens, this approach is optimal for generating complete phylogenetic sampling for clades of interest and relatively cheaply contributing confidently resolved twigs to the Tree of Life.

## Acknowledgements

The UCR seed grant “Unlocking the Vault of SoCal Biota” awarded to CW, Amy Litt, and John Heraty and a Dr. Mir S. Mulla and Lelia Mulla Endowed Scholarship awarded to AK are acknowledged for supporting this project. We would like to thank Randall Schuh (AMNH), Serguei Triapitsyn (UCRC), and Fedor Konstantinov (ZISP) for loaning material and permitting dissections and DNA extractions, Randall Schuh (AMNH) and members of the Weirauch lab for reviewing the manuscript.

## Data Accessibility

-DNA sequences: GenBank accessions [annotated mitochondrial genomes, ribosomal genes, and dynein fragments for baits will be uploaded to GenBank, and accession numbers will be indicated in Table S3]; NCBI SRA: SRP136090, accession numbers for individual samples are indicated in the Table S3.

-Final DNA sequence alignments and partitioning schemes: will be uploaded to Dryad repository.

-Voucher specimen information including photographs: available through the Plant Bug Planetary Biodiversity Inventory Project website (http://research.amnh.org/pbi/heteropteraspeciespage/), linked to the unique specimen identifier (See Table 1, the USI column) [photographs are in the process of being uploaded].

## Author contributions

AK, ERLG & CW designed the research, AK and CW performed the research, AK analyzed the data and AK, ERLG and CW wrote the paper.

## References

Anisimova, M., & Gascuel, O. (2006). Approximate likelihood-ratio test for branches: a fast, accurate, and powerful alternative. Systematic biology, 55(4), 539–552. doi: 10.1080/10635150600755453

Bankevich, A., Nurk, S., Antipov, D., Gurevich, A. A., Dvorkin, M., Kulikov, A. S.,… & Pevzner P. A. (2012). SPAdes: a new genome assembly algorithm and its applications to single-cell sequencing. Journal of computational biology, 19(5), 455–477. doi: 10.1089/cmb.2012.0021

Bi, K., Linderoth, T., Vanderpool, D., Good, J. M., Nielsen, R., & Moritz, C. (2013). Unlocking the vault: next-generation museum population genomics. Molecular ecology, 22 (24), 6018–6032. doi: 10.1111/mec.12516

Blaimer, B. B., Lloyd, M. W., Guillory, W. X., & Brady, S. G. (2016). Sequence capture and phylogenetic utility of genomic ultraconserved elements obtained from pinned insect specimens. PloS one, 11(8), e0161531. doi: 10.1371/journal.pone.0161531

Bolger, A. M., Lohse, M., & Usadel, B. (2014). Trimmomatic: a flexible trimmer for Illumina sequence data. Bioinformatics, 30(15), 2114–2120. doi: 10.1093/bioinformatics/btu170

Burrell, A. S., Disotell, T. R., & Bergey, C. M. (2015). The use of museum specimens with high-throughput DNA sequencers. Journal of human evolution, 79, 35–44. doi: 10.1016/j.jhevol.2014.10.015

Cassis, G., & Schuh, R. T. (2012). Systematics, biodiversity, biogeography, and host associations of the Miridae (Insecta: Hemiptera: Heteroptera: Cimicomorpha). Annual review of entomology, 57, 377–404. doi: 10.1146/annurev-ento-121510-133533

Coddington, J. A., Agnarsson, I., Miller, J. A., Kuntner, M., & Hormiga, G. (2009). Undersampling bias: the null hypothesis for singleton species in tropical arthropod surveys. Journal of animal ecology, 78(3), 573–584. doi: 10.1111/j.1365-2656.2009.01525.x

Cridland, J. M., Ramirez, S. R., Dean, C. A., Sciligo, A., & Tsutsui, N. D. (2018). Genome sequencing of museum specimens reveals rapid changes in the genetic composition of honey bees in California. Genome biology and evolution, 10(2), 458–472. doi: 10.1093/gbe/evy007

Cusimano, N., & Renner, S. S. (2010). Slowdowns in diversification rates from real phylogenies may not be real. Systematic biology, 59(4), 458–464. doi: 10.1093/sysbio/syq032

Cusimano, N., Stadler, T., & Renner, S. S. (2012). A new method for handling missing species in diversification analysis applicable to randomly or nonrandomly sampled phylogenies. Systematic biology, 61(5), 785–792. doi: 10.1093/sysbio/sys031

DiEuliis, D., Johnson, K. R., Morse, S. S., & Schindel, D. E. (2016). Opinion: Specimen collections should have a much bigger role in infectious disease research and response. Proceedings of the National Academy of Sciences, 113(1), 4–7. doi: 10.1073/pnas.1522680112

Fisher, S., Barry, A., Abreu, J., Minie, B., Nolan, J., Delorey, T. M.,… & Berlin, A. M. (2011). A scalable, fully automated process for construction of sequence-ready human exome targeted capture libraries. Genome biology, 12(1), R1. doi: 10.1186/gb-2011-12-1-r1

Glenn, T. C., Nilsen, R., Kieran, T. J., Finger, J. W., Pierson, T. W., Bentley, K. E.,… & Faircloth, B. C. (2016). Adapterama I: universal stubs and primers for thousands of dual-indexed Illumina libraries (iTru & iNext). BioRxiv, 049114. doi: 10.1101/049114

Hallmann, C. A., Sorg, M., Jongejans, E., Siepel, H., Hofland, N., Schwan, H.,… & de Kroon, H. (2017). More than 75 percent decline over 27 years in total flying insect biomass in protected areas. PloS one, 12(10), e0185809. doi: 10.1371/journal.pone.0185809

Hartley, C. J., Newcomb, R. D., Russell, R. J., Yong, C. G., Stevens, J. R., Yeates, D. K.,… & Oakeshott, J. G. (2006). Amplification of DNA from preserved specimens shows blowflies were preadapted for the rapid evolution of insecticide resistance. Proceedings of the National Academy of Sciences, 103(23), 8757–8762. doi: 10.1073/pnas.0509590103

Höhna, S., Stadler, T., Ronquist, F., & Britton, T. (2011). Inferring speciation and extinction rates under different sampling schemes. Molecular biology and evolution, 28(9), 2577–2589. doi: 10.1093/molbev/msr095

Jung, S., & Lee, S. (2011). Molecular phylogeny of the plant bugs (Heteroptera: Miridae) and the evolution of feeding habits. Cladistics, 28(1), 50–79. doi: 10.1111/j.1096-0031.2011.00365.x

Kalyaanamoorthy, S., Minh, B. Q., Wong, T. K., von Haeseler, A., & Jermiin, L. S. (2017). ModelFinder: fast model selection for accurate phylogenetic estimates. Nature methods, 14(6), 587. doi: 10.1038/nmeth.4285

Kanda, K., Pflug, J. M., Sproul, J. S., Dasenko, M. A., & Maddison, D. R. (2015). Successful recovery of nuclear protein-coding genes from small insects in museums using Illumina sequencing. PLoS One, 10(12), e0143929. doi: 10.1371/journal.pone.0143929

Katoh, K., & Standley, D. M. (2013). MAFFT multiple sequence alignment software version 7: improvements in performance and usability. Molecular biology and evolution, 30(4), 772–780. doi: 10.1093/molbev/mst010

Kearse, M., Moir, R., Wilson, A., Stones-Havas, S., Cheung, M., Sturrock, S.,… & Drummond A. (2012). Geneious Basic: an integrated and extendable desktop software platform for the organization and analysis of sequence data. Bioinformatics, 28(12), 1647–1649. 10.1093/bioinformatics/bts199

Konstantinov, F. V., & Knyshov, A. A. (2015). The tribe Bryocorini (Insecta: Heteroptera: Miridae: Bryocorinae): phylogeny, description of a new genus, and adaptive radiation on ferns. Zoological Journal of the Linnean Society, 175(3), 441–472. doi: 10.1111/zoj.12283

Lanfear, R., Frandsen, P. B., Wright, A. M., Senfeld, T., & Calcott, B. (2016). PartitionFinder 2: new methods for selecting partitioned models of evolution for molecular and morphological phylogenetic analyses. Molecular Biology and Evolution, 34(3), 772–773. doi: 10.1093/molbev/msw260

Lee, H. Y., Park, M. J., Kim, N. Y., Sim, J. E., Yang, W. I., & Shin, K. J. (2010). Simple and highly effective DNA extraction methods from old skeletal remains using silica columns. Forensic Science International: Genetics, 4(5), 275–280. doi: 10.1016/j.fsigen.2009.10.014

Li, C., Corrigan, S., Yang, L., Straube, N., Harris, M., Hofreiter, M.,… & Naylor, G. J. (2015). DNA capture reveals transoceanic gene flow in endangered river sharks. Proceedings of the National Academy of Sciences, 112(43), 13302–13307. doi: 10.1073/pnas.1508735112

Li, C., Hofreiter, M., Straube, N., Corrigan, S., & Naylor, G. J. (2013). Capturing protein-coding genes across highly divergent species. Biotechniques, 54(6), 321–326. doi: 10.2144/000114039

Li, H., & Durbin, R. (2009). Fast and accurate short read alignment with Burrows–Wheeler transform. Bioinformatics, 25 (14), 1754–1760. doi: 10.1093/bioinformatics/btp324

Lim, G. S., Balke, M., & Meier, R. (2011). Determining species boundaries in a world full of rarity: singletons, species delimitation methods. Systematic biology, 61(1), 165–169. doi: 10.1093/sysbio/syr030

Liu, S., Wang, X., Xie, L., Tan, M., Li, Z., Su, X.,… & Zhou, X. (2016). Mitochondrial capture enriches mito-DNA 100 fold, enabling PCR-free mitogenomics biodiversity analysis. Molecular ecology resources, 16(2), 470–479. doi: 10.1111/1755-0998.12472

Maddison, D. R., & Cooper, K. W. (2014). Species delimitation in the ground beetle subgenus Liocosmius (Coleoptera: Carabidae: Bembidion), including standard and next-generation sequencing of museum specimens. Zoological Journal of the Linnean Society, 172(4), 741–770. doi: 10.1111/zoj.12188

Mariac, C., Scarcelli, N., Pouzadou, J., Barnaud, A., Billot, C., Faye, A.,… & Couvreur, T. L. P. (2014). Cost-effective enrichment hybridization capture of chloroplast genomes at deep multiplexing levels for population genetics and phylogeography studies. Molecular Ecology Resources, 14(6), 1103–1113. doi: 10.1111/1755-0998.12258

Maricic, T., Whitten, M., & Pääbo, S. (2010). Multiplexed DNA sequence capture of mitochondrial genomes using PCR products. PloS one, 5(11), e14004. doi: 10.1371/journal.pone.0014004

McCormack, J. E., Tsai, W. L., & Faircloth, B. C. (2015). Sequence capture of ultraconserved elements from bird museum specimens. Molecular ecology resources, 16(5), 1189–1203. doi: 10.1111/1755-0998.12466

Menard, K. L., Schuh, R. T., & Woolley, J. B. (2014). Total-evidence phylogenetic analysis and reclassification of the Phylinae (Insecta: Heteroptera: Miridae), with the recognition of new tribes and subtribes and a redefinition of Phylini. Cladistics, 30(4), 391–427. doi: 10.1111/cla.12052

Minh, B. Q., Nguyen, M. A. T., & von Haeseler, A. (2013). Ultrafast approximation for phylogenetic bootstrap. Molecular biology and evolution, 30(5), 1188–1195. doi: 10.1093/molbev/mst024

Mitchell, A. (2015). Collecting in collections: a PCR strategy and primer set for DNA barcoding of decades-old dried museum specimens. Molecular ecology resources, 15 (5), 1102–1111. doi: 10.1111/1755-0998.12380

Namyatova, A. A., Konstantinov, F. V., & Cassis, G. (2015). Phylogeny and systematics of the subfamily Bryocorinae with the emphasis on the tribe Dicyphini sensu Schuh, 1976 derived from morphological characters. Systematic Entomology, 41, 3–40. doi: 10.1111/syen.12140

Nguyen, L. T., Schmidt, H. A., von Haeseler, A., & Minh, B. Q. (2014). IQ-TREE: a fast and effective stochastic algorithm for estimating maximum-likelihood phylogenies. Molecular biology and evolution, 32(1), 268–274. doi: 10.1093/molbev/msu300

Paradis, E., Claude, J., & Strimmer, K. (2004). APE: analyses of phylogenetics and evolution in R language. Bioinformatics, 20(2), 289–290. doi: 10.1093/bioinformatics/btg412

Peñalba, J. V., Smith, L. L., Tonione, M. A., Sass, C., Hykin, S. M., Skipwith, P. L.,… & Moritz, C. (2014). Sequence capture using PCR-generated probes: a cost-effective method of targeted high-throughput sequencing for nonmodel organisms. Molecular Ecology Resources, 14(5), 1000–1010. doi: 10.1111/1755-0998.12249

Revell, L. J. (2012). phytools: an R package for phylogenetic comparative biology (and other things). Methods in Ecology and Evolution, 3(2), 217–223. doi: 10.1111/j.2041-210X.2011.00169.x

Rohland, N., & Reich, D. (2012). Cost-effective, high-throughput DNA sequencing libraries for multiplexed target capture. Genome research, 22(5), 939–946. doi: 10.1101/gr.128124.111

Schmieder, R., & Edwards, R. (2011). Quality control and preprocessing of metagenomic datasets. Bioinformatics, 27(6), 863–864. doi: 10.1093/bioinformatics/btr026

Schuh, R. T. (2004). Revision of Tuxedo Schuh (Hemiptera: Miridae: Phylinae). American Museum Novitates, 3435, 1–26. doi: 10.1206/0003-0082(2004)435<0001:ROTSHM>2.0.CO;2

Staats, M., Erkens, R. H., van de Vossenberg, B., Wieringa, J. J., Kraaijeveld, K., Stielow, B.,… & Bakker, F. T. (2013). Genomic treasure troves: complete genome sequencing of herbarium and insect museum specimens. PLoS One, 8 (7), e69189.

Stamatakis, A. (2014). RAxML version 8: a tool for phylogenetic analysis and post-analysis of large phylogenies. Bioinformatics, 30(9), 1312–1313. doi: 10.1093/bioinformatics/btu033

Stamatakis, A., Hoover, P., & Rougemont, J. (2008). A rapid bootstrap algorithm for the RAxML web servers. Systematic biology, 57(5), 758–771. doi: 10.1080/10635150802429642

Stork, N. E. (2018). How Many Species of Insects and Other Terrestrial Arthropods Are There on Earth?. Annual review of entomology, 63. doi: 10.1146/annurev-ento-020117-043348

Stork, N. E., McBroom, J., Gely, C., & Hamilton, A. J. (2015). New approaches narrow global species estimates for beetles, insects, and terrestrial arthropods. Proceedings of the National Academy of Sciences, 112(24), 7519–7523. doi: 10.1073/pnas.1502408112

Suchan, T., Pitteloud, C., Gerasimova, N. S., Kostikova, A., Schmid, S., Arrigo, N.,… & Alvarez, N. (2016). Hybridization capture using RAD probes (hyRAD), a new tool for performing genomic analyses on collection specimens. PloS one, 11(3), e0151651. doi: 10.1371/journal.pone.0151651

Tatarnic, N. J., & Cassis, G. (2012). The Halticini of the world (Insecta: Heteroptera: Miridae: Orthotylinae): generic reclassification, phylogeny, and host plant associations. Zoological Journal of the Linnean Society, 164(3), 558–658. doi: 10.1111/j.1096-3642.2011.00770.x

Weirauch, C. (2006a). New genera, new species, and new combinations in western Nearctic Phylini (Heteroptera: Miridae: Phylinae). American Museum Novitates, 3521, 1–41. doi: 10.1206/0003-0082(2006)3521[1:NGNSAN]2.0.CO;2

Weirauch, C. (2006b). New genera and species of oak-associated Phylini (Heteroptera: Miridae: Phylinae) from western North America. American Museum Novitates, 3522, 1–54. doi: 10.1206/0003-0082(2006)3522[1:NGASOO]2.0.CO;2

Weirauch, C., Seltmann, K. C., Schuh, R. T., Schwartz, M. D., Johnson, C., Feist, M. A., & Soltis, P. S. (2016). Areas of endemism in the Nearctic: a case study of 1339 species of Miridae (Insecta: Hemiptera) and their plant hosts. Cladistics, 33(3), 279–294. doi: 10.1111/cla.12169

Yang, D. Y., Eng, B., Waye, J. S., Dudar, J. C., & Saunders, S. R. (1998). Improved DNA extraction from ancient bones using silica-based spin columns. American journal of physical anthropology, 105(4), 539–543. doi: 10.1002/(SICI)1096-8644(199804)105:4<539::AID-AJPA10>3.0.CO;2-1

Yin, T., Cook, D., & Lawrence, M. (2012). ggbio: an R package for extending the grammar of graphics for genomic data. Genome biology, 13(8), R77. doi: 10.1186/gb-2012-13-8-r77

Yu, G., Smith, D. K., Zhu, H., Guan, Y., & Lam, T. T. Y. (2016). ggtree: an R package for visualization and annotation of phylogenetic trees with their covariates and other associated data. Methods in Ecology and Evolution, 8(1), 28–36. doi: 10.1111/2041-210X.12628

